# Optical tuning of polymer functionalized zinc oxide quantum dots as a selective probe for specific detection of antibiotics

**DOI:** 10.1101/2022.11.04.515220

**Authors:** Awadhesh Kumar Verma, Tarun Kumar Dhiman, GBVS Lakshmi, Pratima Solanki

## Abstract

It is crucial to monitor the antibiotic levels in the environment like water, food products in the current scenario as the antibiotic concentration above threshold is harmful for the human health. Excess consumption of antibiotics leads to antibiotic resistance that hinders the control and cure of microbial diseases. So, these challenges motivated to devise an optical nano-senor which can sense the ultra-low concentration of antibiotics. In this proposed research work, emphasis is to develop a method which is simple and selective to analyze the detection and presence of antibiotics in various samples like tape water milk etc. using fluorescent ZnO QDs based nano-sensor. For this, fluorescent and different polymers (polyvinylalcohol – PVA and polyvinylpyrrolidine – PVP) capped ZnO QDs were synthesized using modified sol-gel technique. These were used as fluorescent probe to monitor the presence of antibiotics. The optical characterizations of synthesized QDs were performed using UV-Visible absorption & fluorescence spectroscopic methods while structural characteristics were analyzed by using FTIR, XRD and EDX. Charge on the synthesized QDs were obtained with the help of ZETA potential. Here ten different antibiotics were targeted among, Ciprofloxacin and Moxifloxacin have shown excellent sensing and specificity with PVA-ZnO QDs and PVP-ZnO QDs respectively.

## 1. Introduction

Antibiotic, a corner stone of modern medicinal industry is the drug or medicine which stops or kills the growth of microbial diseases. The term antibiotic [1] derived from word “antibiosis” in 1889 by Louis Velillemin a student of Louis Pasteur. The literal meaning of antibiotic is Anti-means against and bios means-life i.e., anything against life or can be used to destroy the life. Simply, antibiotics are the chemical substances produced by various microorganisms in low concentrations which destroy or kills or inhibit the growth of other microorganisms. Antibiotics may have either “broad” or “narrow” spectrum. Bacteria fall under the “broad” spectrum are agile against numerous divergent category of bacteria, while in “narrow” spectrum it is agile against very little varieties of bacteria[2]. The emergence of antibiotics for the therapeutics of several contagious diseases in human beings including diverse category of animals has been remarked as the inception of antimicrobial therapeutic era in 1940s. These are mostly used due to their particular activity to counteract both gram positive as well as gram negative bacteria. In Global scenario, India, China and USA are in top three countries for the highest antibiotics consumption. From the past data analysis, it is clear that the countries having higher per capita consumption of antibiotics have higher antibiotics resistance rate.[3], [4] The incompetence for diagnosing bactericidal infections is the major constrain that may lead to indecorous use of antibiotics, also there is a reduction in rate of survival in septic situations [5]. Figure 1 shows the antibiotic resistance mechanism.

**Fig. 1:**
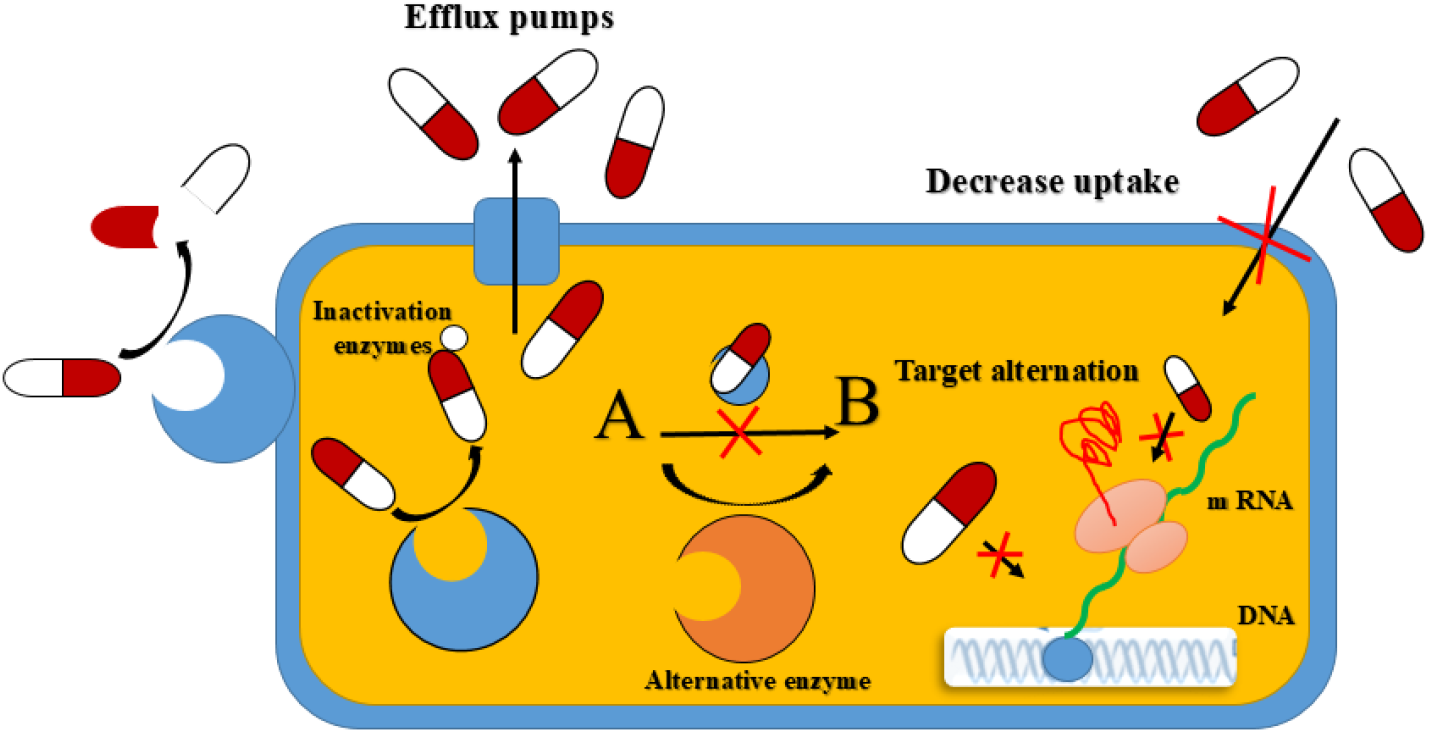
Schematic showing the mechanism of antibiotic resistance bacteria [6]

There is a global concern that antibiotics are vigorously contaminating our environment. Antibiotics has been detected in limited proportions in ground water. The accurate detection of antibiotics level in biological and clinical specimen is an urgent need due to its potential toxic effects. Till now the detection of antibiotics was performed by conventional approach such as high-performance liquid chromatography (HPLC),[7] fluorometry[8], spectrophotometry [9] etc. Although HPLC is widely used owing to its capability, susceptibility and selectivity, it is highly time consuming and needs large amount of solvent usage. Also, these high-level devices require higher cost of maintenance which narrow down its applicability. Electrochemical technique for the detection of analyte based on nanomaterials electrodes modified with nanomaterial is a meticulous approach used in analytical chemistry to determine drugs as well as biomolecules.[10] Few immunosensors for the detection of various antibiotics were reported in the literature [references]. But the main advantage of optical sensing over the electrochemical immunosensors is that, in optical sensors, no other biomolecules like aptamer, antibodies etc. are required. The targeted molecule can be detected directly based on the specificity of the nanomaterials and hence reduces the cost and time. As no biomolecule is used, they can be operated at ambient temperatures as well as the humid environment. Apart from the above features this type of sensors are fast reliable and reproducible.[11]–[16]

Enormous reports have been published regarding monodispersed, stable and fluorescent ZnO QDs in several organic media[17]. ZnO QDs being highly active sometimes undergo agglomeration which leads to poor dispersion and loses all their fluorescence. Several methods and techniques are therefore, established to overcome the challenges such as modification of surface of ZnO nanomaterials making use of water soluble/organic ligands, silanes or by capping of ZnO nanomaterials with polymer. Prachi Joshi et al. have gone through synthesis of highly fluorescent polyethyleneimine (PEI) capped ZnO[18]. The fluorescence spectra showed defect related emission around 555 nm in water as dispersion medium and no agglomeration of ZnO showing long range degradation in luminescence. It was evident that surface modification may produce excellent binding affinity for several molecules like nucleic acid, monosaccharide and protein etc., which can make the ZnO QDs as an excellent candidate as fluorescence contrast agent for bioimaging, bioconjugation and other applications.[19], [20]

In this study, PVA and PVP polymers were used as the capping agents for the tuning fluorescence properties of ZnO quantum dots. These polymers as a capping agent imparts charges on quantum dots which enhances the specificity towards certain analytes. Poly vinyl alcohol (PVA) is a synthetic polymer soluble in water. It provides large number of active -OH groups. It forms polymer complex of metal ion through ligand interaction. ZnO nanomaterials synthesized without using these polymers show very less emission. But PVA functionalized ZnO QDs have *Zn*+ defects which play a major role in enhancing the fluorescence intensity. [21]–[23] Polyvinylpyrrolidone (PVP) is a water-soluble polymer, and commonly used as viscosity modifier in nano field. Water dispersible ZnO QDs have been synthesized using sol–gel method as the highly fluorescent nanomaterial in which water acts as a solvent while PVP act as a viscosity modifying agent. This helps to enhance fluorescence intensity of the quantum dots hence we can make the highly fluorescent quantum dots with tuned optical properties.[24], [25]

In this proposed research work, emphasis is to develop a method which is simple and selective to the detection of antibiotics in various sample like tape water milk etc. using fluorescent ZnO QDs based nano-sensor. For this, fluorescent and polymer capped ZnO QDs using modified solgel technique were synthesized. Two polymers PVA and PVP were used as capping and optical tuning agents for ZnO QD_S_. These were used as fluorescent probes to detect two different antibiotics selectively. The PVA capped ZnO QDs (PVA-ZnO QDs) were used in the detection of Ciprofloxacin and PVP capped ZnO QDs (PVP-ZnO QDs) were used for Moxifloxacin detection.

## 2. Experimental Section

### 2.1 Materials

Polyvinyl Alcohol PVA (Cold) was purchased form Central Drug House (P) Ltd, India. Polyvinylpyrrolidone (PVP K-30) pure was purchased from Sisco Research Laboratories Pvt. Ltd., India. Zinc acetate dihydrate, Moxifloxacin hydrochloride (C_21_H_24_FN_3_O_4_), citric acid (CA) (C_6_H_8_O_7_), cysteine (Cys)(C_3_H_7_NO_2_S) and bovine serum albumin (BSA) were procured from Sigma-Aldrich. Uric acid (UA) (C_5_H_4_N_4_O_3_), sodium hydroxide pellets (NaOH), cholesterol (Cho) (C_27_H_46_O), norfloxacin (Nor) (C_16_H_18_FN_3_O_3_), levofloxacin (Levo) (C_18_H_20_FN_3_O_4_), ofloxacin (Oflo) (C_18_H_20_FN_3_O4), ciprofloxacin hydrochloride (Cipro) (C_17_H_18_FN_3_O_4_·HCl), gentamicin sulfate (Genta) (C_21_H_43_N_5_O_7_·H_2_SO_4_), florfenicol (Flor) (C_12_H_14_C_l2_FNO_4_S), erythromycin (Erythro) (C_37_H_67_NO_13_), thiamphenicol (Thiam) (C_12_H_15_C_l2_NO_5_S), arginine (Arg) (C_6_H_14_N4O_2_), ascorbic acid (AA) (C_6_H_8_O_6_), methionine (Met) (C_5_H_11_NO_2_S), casein (C_81_H_125_N_22_O_39_P), sulfuric acid (H_2_SO_4_) and urea (CH_4_N_2_O) were bought from Sisco Research Laboratories Pvt. Ltd., India. Methanol was procured from Emsure. Folic acid (FA) (C_19_H_19_N_2_O_6_) and aspartic acid (Asp) (NH_2_CH (COOH) CH_2_·COOH) were purchased from CDH. Magnesium chloride (MgC_l2_) was acquired from Thomas Baker.), Tetracycline hydrochloride (Tet) (C_22_H_24_N_2_O_6_·HCl), Ampicillin sodium salt (Amphi) (C_16_H_18_N_3_O_4_SNa) copper chloride (CuCl_2_), and ferrous chloride (FeCl2·H2O) were obtained from Hi-media Laboratory Pvt. Ltd. Glucose (Glu) (C_6_H_12_O_6_) and Sodium chloride (NaCl), were bought from Fine Chem Limited (S.D.F.C.L). Calcium chloride (CaCl_2_) was procured from Rankem Avantor India pvt. Ltd. Potassium chloride was purchased form from Thermo Fisher Scientific Pvt. Ltd. All other chemicals and solvents used are of analytical grade and has been used directly without any further purification. All the solutions were prepared in deionized (DI) water.

### 2.2. Nanomaterial Synthesis

ZnO QDs, PVA-ZnO QDs and PVP-ZnO QDs were separately synthesized using modified solgel approach [55], which involves mixing of precursor solutions (5 mM solution of zinc acetate dihydrates and 10 mM solution of sodium hydroxide prepared separately in Methanol and DI water), refluxing at desired temperature, neutralization of pH followed by characterization. Fig. 2 shows the flow chart of reflexing sol-gel approach for the synthesis of ZnO QDs. Refluxing was performed at 65°C [56]. For heating, a semi spherical type of electrically powered heating mantel was used with a magnetic stirrer to provide constant stirring during heating. For the condensation of vapors, chilled water was circulated in the condenser, while the temperature was measured by placing a thermometer in one of the side necks. The process continued for certain time to ascertain the growth mechanism followed by cooling the solution at room temperature. After cooling, solution pH was neutralized by dropwise adding NaOH solution under continuous stirring until pH 7.0 was obtained. The solution was further stirred for some time to get a uniform pH. The reaction followed during formation of ZnO QDs are as follows:

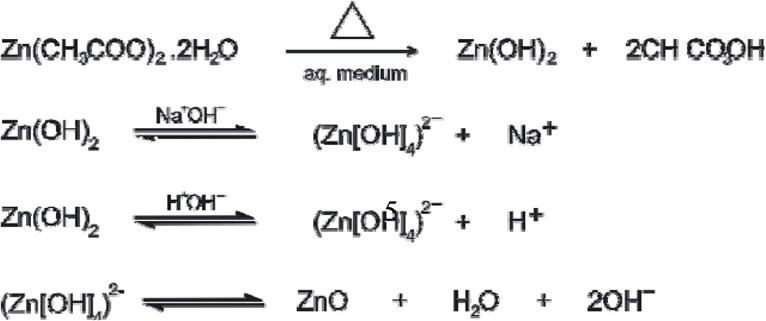

**Fig. 2.**
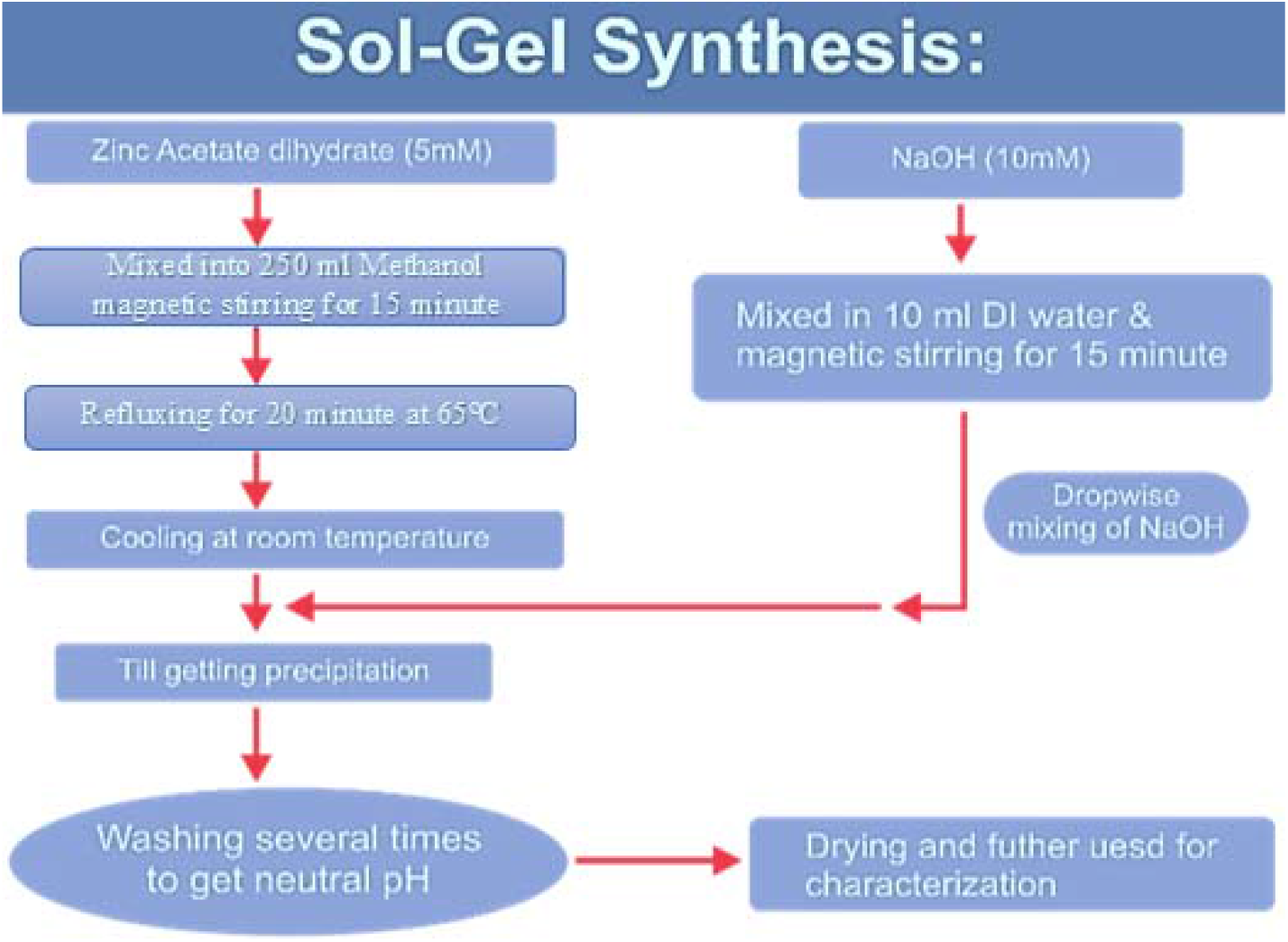
Flow chart showing the details of sol-gel synthesis of ZnO QDs

### 2.3. PVA-ZnO QDs

10 ml of PVA (50 mg/10 ml) solution was mixed drop wise to ZnO-NaOH reaction mixture at room temperature during continuous stirring. The pH of the solution was neutral. The QDs prepared with PVA is abbreviated as PVA -ZnO QDs.

### 2.4. PVP-ZnO QDs

10 ml of PVP solution (50 mg/10 ml) was added as mentioned above and the QDs prepared with PVP is abbreviated as PVP-ZnO QDs.

### 2.5. Nanomaterial Characterization and sensing of the antibiotics

UV-Vis absorption spectroscopy of the as synthesized QDs was carried out in the wavelength range of 190-900 nm using double beam Hitachi (U3900) spectrophotometer. A quartz cuvette of 10 mm path length having fixed slit width (5 nm) and scan rate 600 nm/minute were used for the measurements. Peak absorption wavelength was used as excitation energy for fluorescence measurement. The optical band gap of synthesized ZnO QDs were estimated using Tauc plots [58]. Fluorescence measurements were performed using Cary Eclipse fluorescence spectrophotometer (Agilent Technology, Model G9800A) at a constant excitation and emission slit width of 5 nm and scanning speed of 600 nm/minute. A 3.5 ml quartz cuvette with 10 mm path length was used for measurement. To account the changes in the functional groups, present on the surface of bio-conjugates, FTIR spectroscopy were performed in ATR mode using PerkinElmer Spectrum 1 configuration of FTIR spectrometer unit. The spectra were obtained in the range of 4000 to 400 wavenumber range. X-ray diffraction (XRD) patterns of ZnO QDs, PVA-ZnO QDs and PVP-ZnO QDs were obtained using Ultima IV (Rigaku). Cu-Kα radiation (λ=1.5415 Å) was used as an x-ray source at applied voltage of 40 keV and current 30 mA. The beam incident angle of 60º was kept constant. The sample was made by making a uniform film of ZnO QDs, PVP-ZnO QDs and PVA-ZnO QDs synthesized powder on the sample holder of size 1 cm x 1 cm. The spectra were obtained for the Bragg angle ranging from 10-90° at the scan speed of 8°/min. To determine surface charge and the zeta potential on the quantum dots surface somewhere in diffuse layer in the suspension, we used the Zeta potential analyzer (ZEECOM, Microtech Co. Ltd.). Raman spectroscopy was carried out to study the structural properties of the ZnO QDs, PVA-ZnO QDs and PVP-ZnO QDs. For this we have used Raman-AFM Microscope alpha 300 RA (Oxford Instruments). The time-resolved fluorescence measurement of PVA Capped ZnO and PVP capped ZnO in the absence and presence of antibiotics was recorded at 360 nm emission wavelength using Time Resolved Fluorescence Spectrometer (TRFS) - Edinburgh FL920 Fluorescence Life Time Spectrometer to calculate the average lifetime and quenching effect (static/dynamic). The elemental analysis (EDS) was done by JSM -IT 200.

### 2.6. Sensing

All the nano-conjugates were prepared by physical mixing at room temperature. For sensing of Ciprofloxacin, the concentration range of ciprofloxacin from 1 nM to 1 mM every time mixed with PVA-ZnO solution and PL was measured at every concentration. For sensing of Moxifloxacin, the concentration range of Moxifloxacin from 1 nM to 1 mM every time mixed with PVP-ZnO solution and the PL was measured at every concentration range. For spike study in tap water both Ciprofloxacin and Moxifloxacin were taken. For interference study, different antibiotics along with other interferents were carried out with common interferents such as citric acid (10 mM), ascorbic acid (10 mM), cholesterol (10 mM), aspartic acid (10 mM), uric acid (10 mM), folic acid (10 mM), casein (10 mM), urea (10 mM), glucose (5 mM), sodium (Na^+^), potassium (K^2+^), calcium (Ca^2+^), copper (Cu^2+^), magnesium (Mg^2+^), manganese (Mn^2+^), and Zinc (Zn^2+^) (all ionic solutions 1mM).

## 3. Results and Discussion

Fig. 3 (a) shows optical absorption spectra of as synthesized ZnO QDs (b) PVA-ZnO QDs and (c) PVP-ZnO QDs depicting the absorption peak at around 355 nm, 273 nm and 265 nm, which are in well agreement with reported values for ZnO QDs obtained through sol-gel method [26]. The other peaks in PVA-ZnO and PVP-ZnO are due to the PVA and PVP, respectively. The Tauc plots of ZnO QDs, PVA-ZnO QDs and PVP-ZnO QDs and shown in the respective insets of figures 3(a), 3(b) and 3(c) and the optical bandgap values were obtained approximately as 3 eV, 2.65 eV and 2.57 eV respectively. It is clear from the absorption spectra that the polymer capping of QDs play an important role in tuning the optical absorption and also the optical bandgap. The specific functional groups introduced by capping can further play an important role in the selectivity towards the interaction with antibiotics.

**Fig.3:**
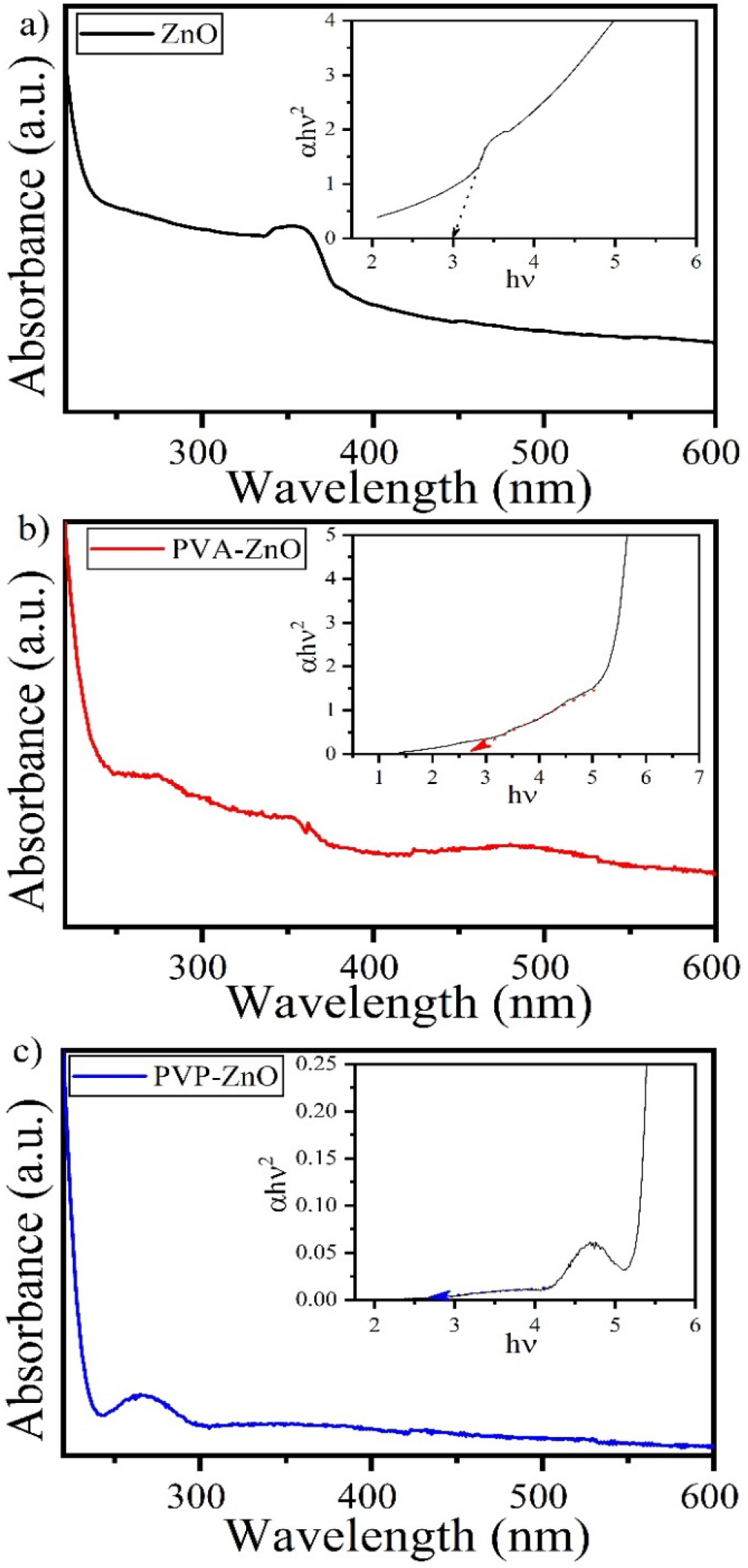
UV-Vis absorbance spectra of ZnO QDs (a), PVA-ZnO QDs and PVP-ZnO QDs respectively. (inset: Corresponding Tauc plots)

Fig.4 shows the fluorescence spectra of as synthesized ZnO QDs, PVA-ZnO QDs and PVP-ZnO QDs obtained at room temperature in the range 250-800 nm with excitation energy of 230 nm. The spectra reveal four emission peaks; the main peak centered around 350 nm, secondary emission peak around 460 nm, a small peak centered at 500 nm and a broad peak between 550-700 nm in agreement with a report for sol-gel synthesized ZnO QDs [27]. Slight left shift in the peaks position has been obtained for the PVA-ZnO QDs and PVP-ZnO QDs in accordance with the optical bandgap variation after polymer capping. The optical parameters obtained are summarized in table 1.

**Table 1.**
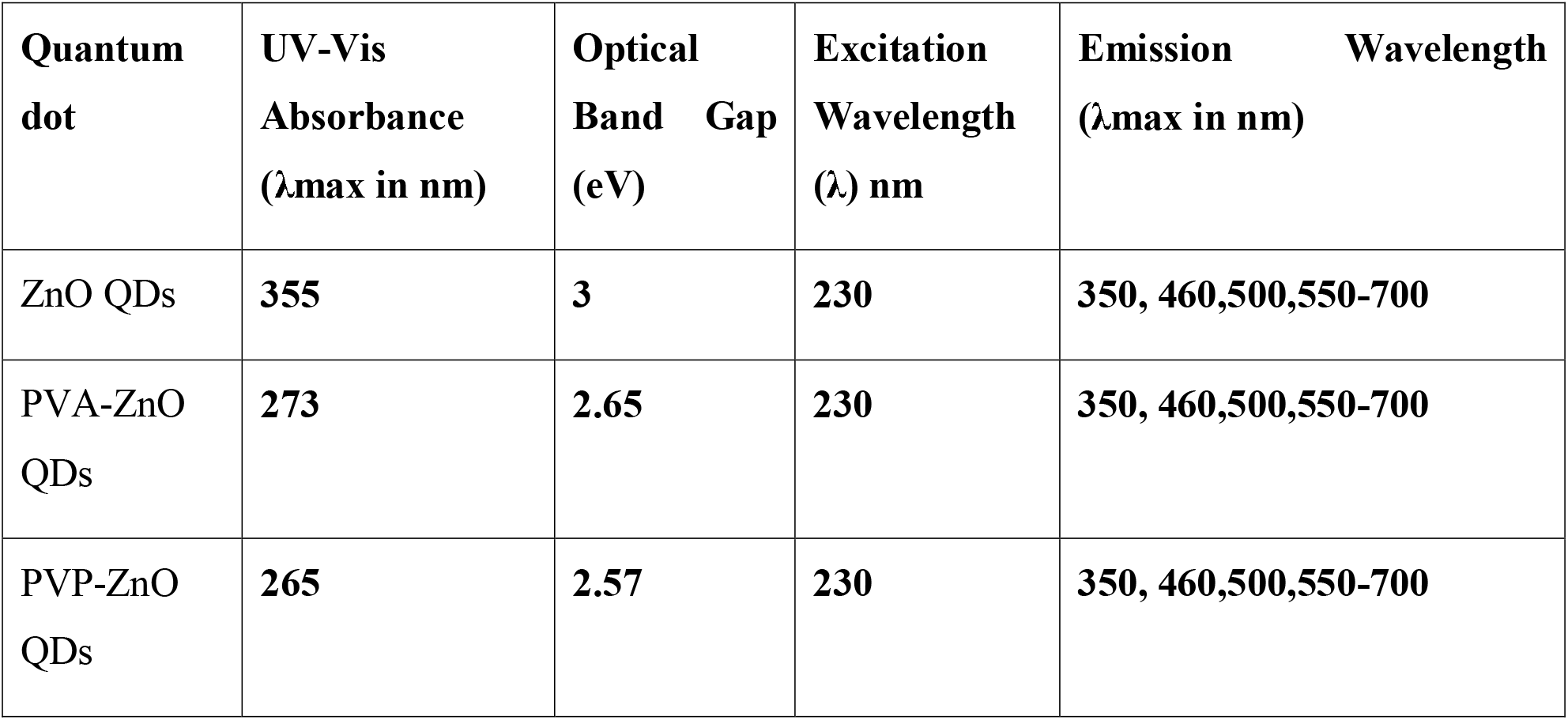
the UV–Vis absorption maxima, optical band gap, excitation wavelength and emission wavelength of synthesized (ZnO QDs, and PVA-ZnO QDs and PVP-ZnO QDs)

**Fig.4:**
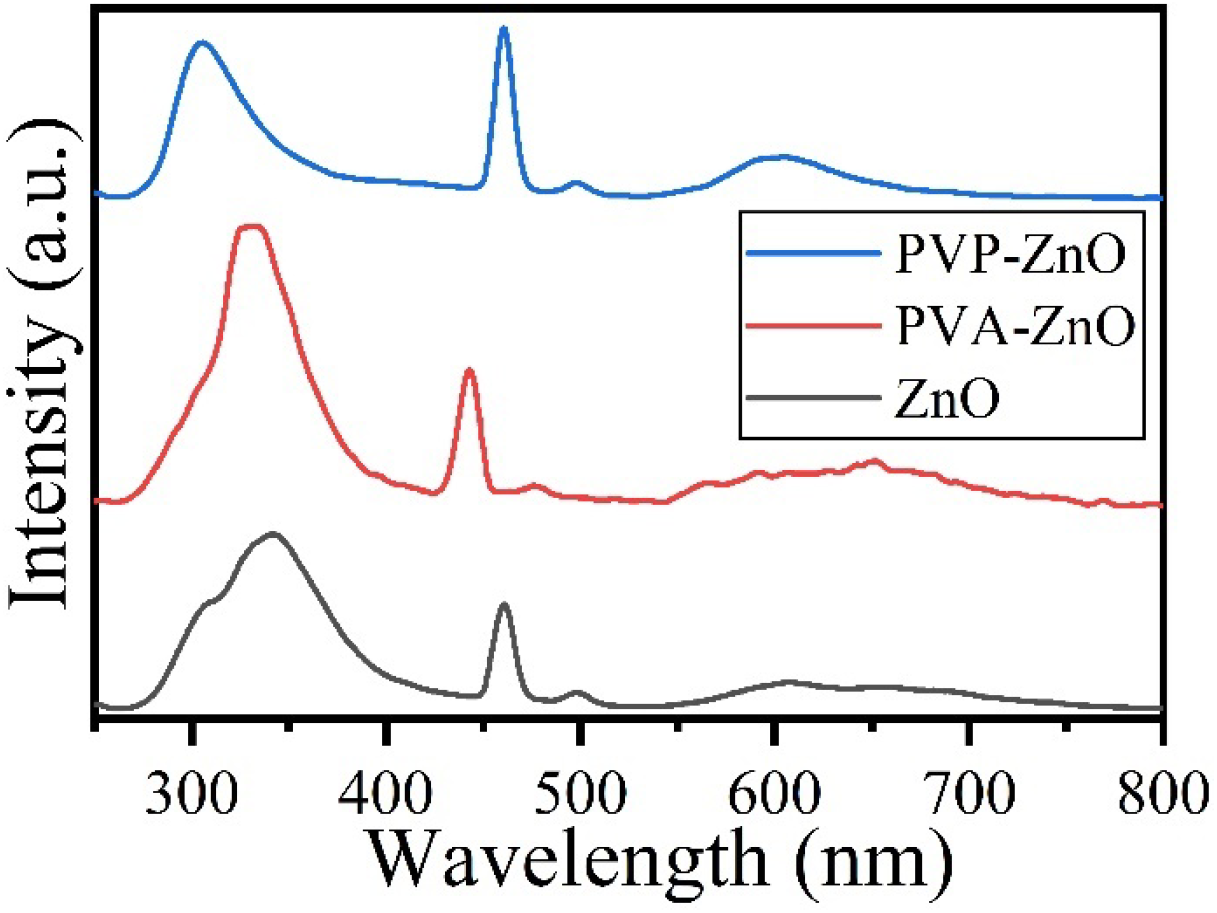
Fluorescence spectrum of ZnO QDs, PVA-ZnO QDs and PVP-ZnO QDs

Fig.5 depicts the FTIR spectrum of as synthesized ZnO QDs, PVP-ZnO QDs and PVA-ZnO QDs. FTIR spectra has been obtained approx. 1014 cm^-1^ which confirms the synthesis of metal oxide quantum dots.

**Fig.5:**
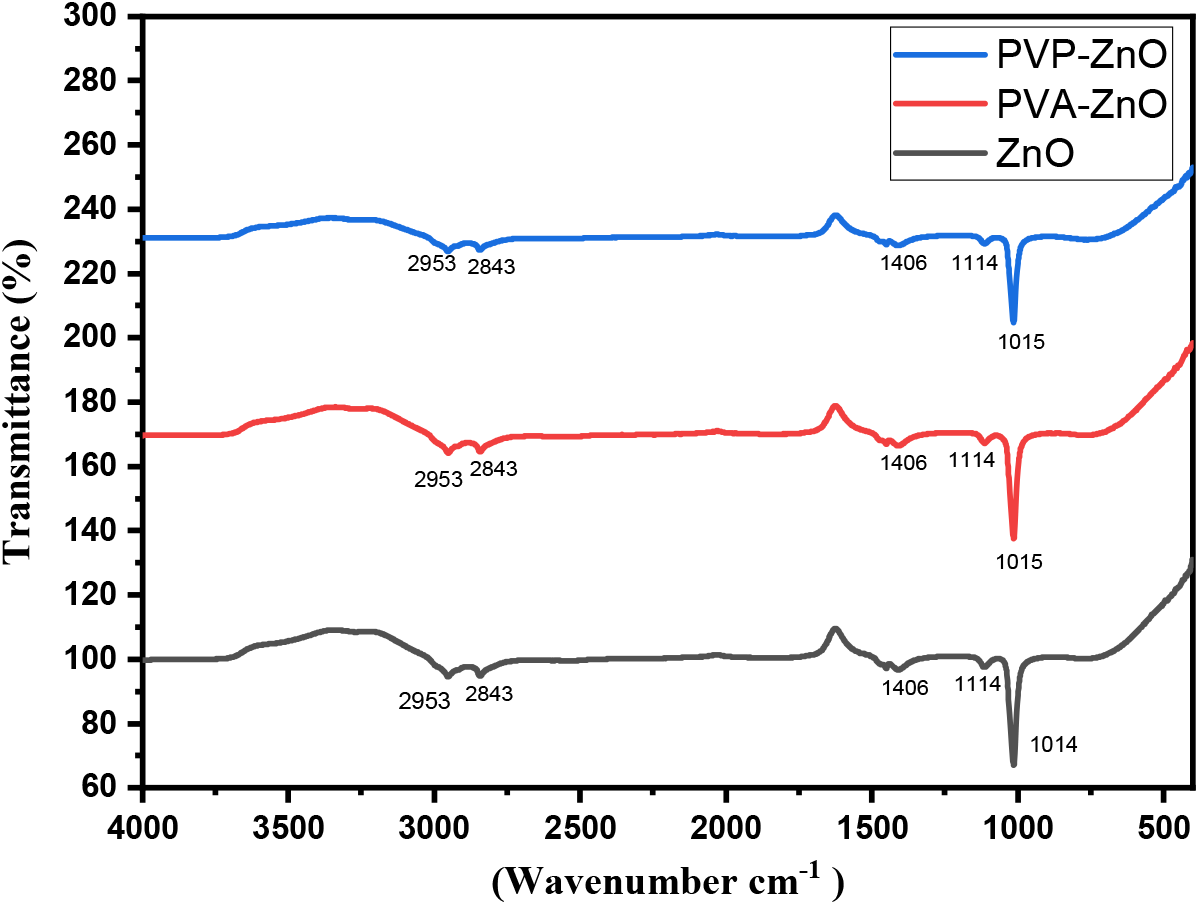
FTIR spectrum of ZnO QDs, PVA-ZnO QDs and PVP-ZnO QDs.

The XRD was performed to study the presence of any impurity in the prepared samples. The three samples were drop casted on glass slides and allowed to dry in air for 24 hours. Same process was repeated 9 more times to obtain a thick layer of ZnO QDs, PVA-ZnO QDs, and PVP-ZnO QDs. No crystalline structure can be seen in the XRD spectra of each sample confirming the purity of QDs, as it is known that quantum dots do not show any sharp peaks in XRD spectra. The broad humps shown in Fig.6 confirmed the small size of the QDs.

**Fig.6:**
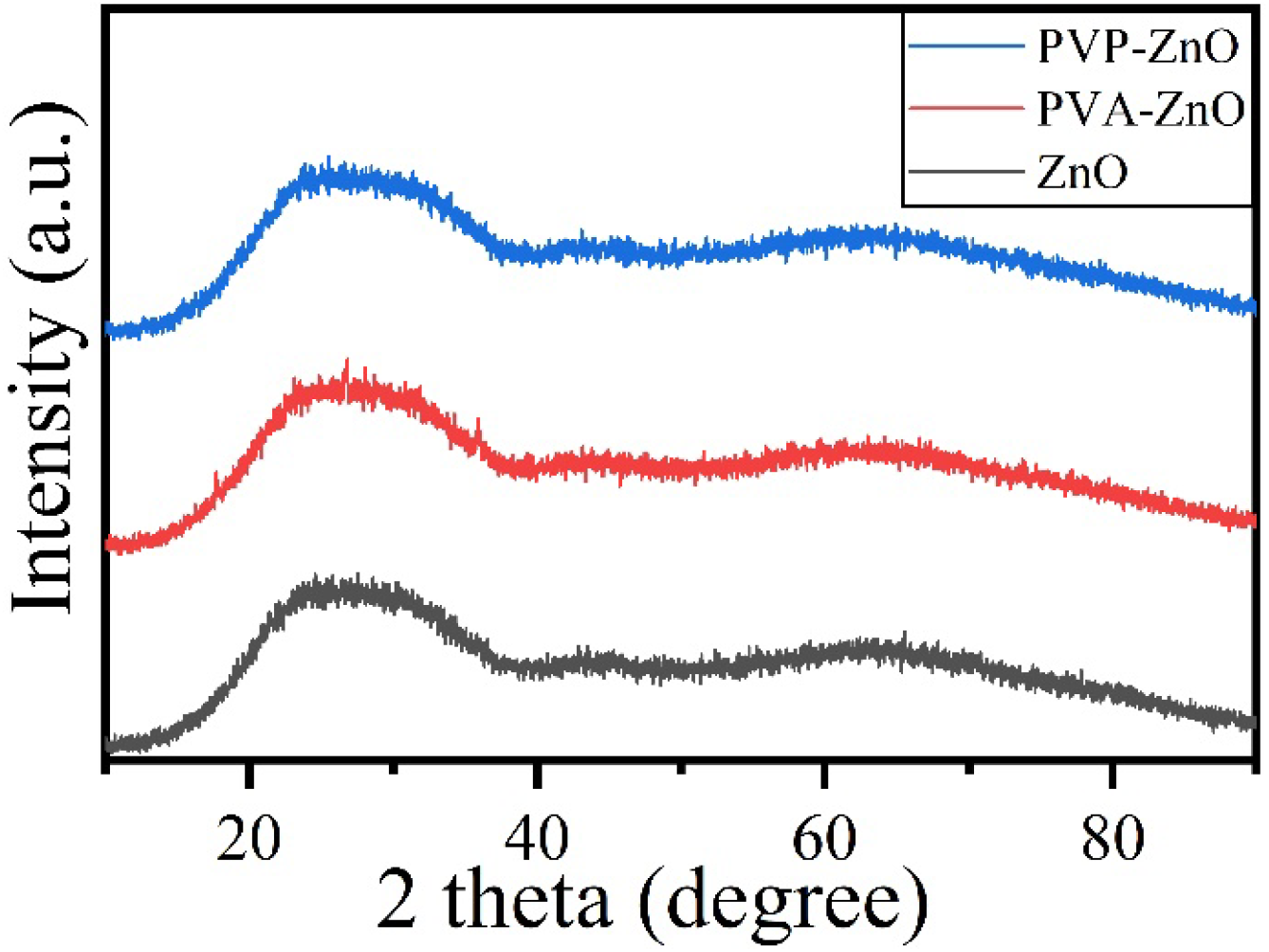
XRD pattern of ZnO QDs, PVA-ZnO QDs and PVP-ZnO QDs

As the QDs have sizes less than 10 nm, the SEM images did not show any specific informative results. Whereas, the energy dispersive x-ray spectroscopy (EDS) was performed to find the elemental composition of the prepared samples. In each of the samples, carbon, oxygen, zinc and sodium were obtained as shown in figure 7 (a), (b) and (c). Carbon is due to the carbon tape used for EDS measurement, while Na is due to the use of NaOH while synthesizing the samples. The presence of Zinc and Oxygen confirms the formation of ZnO quantum dots in all the samples as shown in Fig.7.

**Fig. 7:**
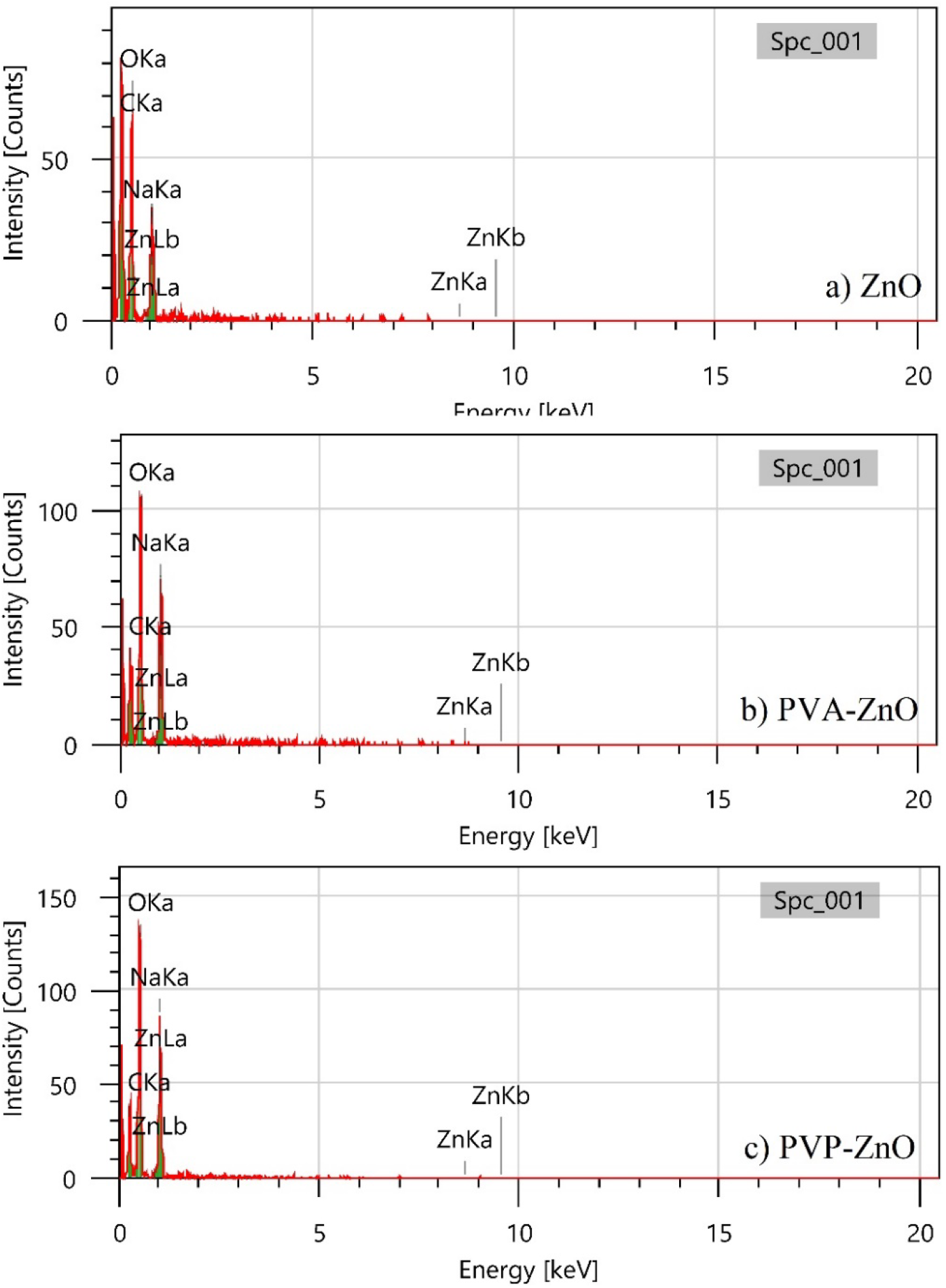
EDS spectra of a) ZnO QDs, b) PVA-ZnO QDs, and c) PVP-ZnO QDs.

Raman Spectroscopy of ZnO QDs, PVA-ZnO QDs and PVP-ZnO QDs were carried out and the spectra is shown in figure 8. There are five common peaks corresponding to ZnO have been obtained at wavenumbers of 369.59, 481.95, 929.35, and 1095.15 cm^−1^ in the Raman spectra of each sample. The peak at 369.59 cm-1 is due to the E2_H_ peak, while the small peak at 481.95 cm^-1^ is due to the A1(LO) mode. The peak at 929.35 cm^−1^ corresponds to A1(TO)+E2_L_. The peak observed at 1095.15 cm^-1^ is due to the 2 (LO) mode. The rest of the peaks in PVA-ZnO QDs and PVP-ZnO QDs are due to the PVA and PVP, respectively.

**Fig. 8:**
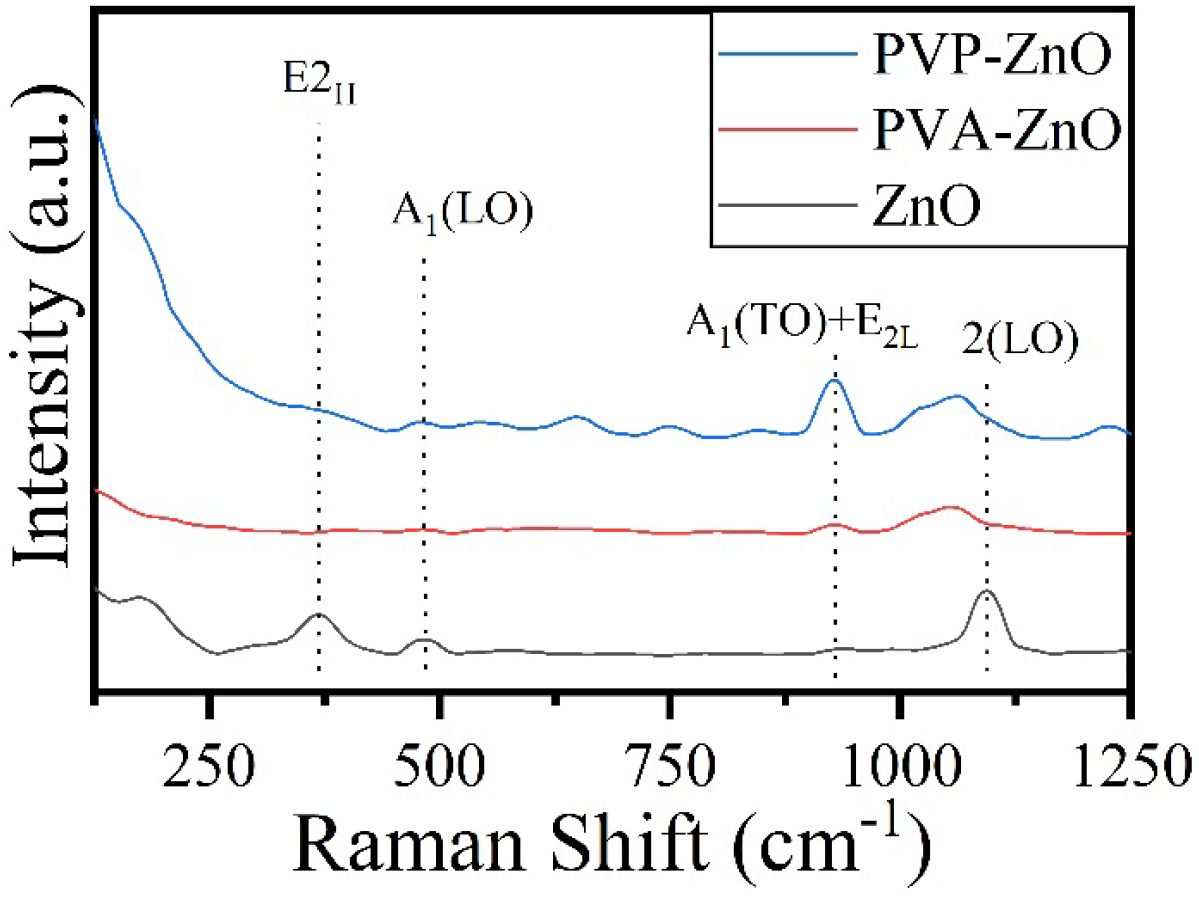
Raman spectra of a) ZnO QDs, b) PVA-ZnO QDs, and c) PVP-ZnO QDs.

Surface charge plays a critical role in surface properties of the nanomaterials. Zeta potential study helps in studying the surface charge of the nanoparticle. Here zeta study has been performed on ZnO QDs, PVP-ZnO QDs and PVA-ZnO QDs. Histogram has been plotted for each sample as shown in figure 9. In each of the sample positive charge is obtained varying in the value. ZnO QDs has zeta potential ranging from 20-180 mV with maximum zeta potential centered at 100-110 mV. PVA-ZnO QDs has zeta potential ranging from 100-300 mV with maximum zeta potential centered at 150-250 mV. PVP-ZnO QDs has zeta potential ranging from 100-350 mV with maximum zeta potential centered at 200-300 mV. The zeta potential increased with capping of the polymers.

**Fig. 9:**
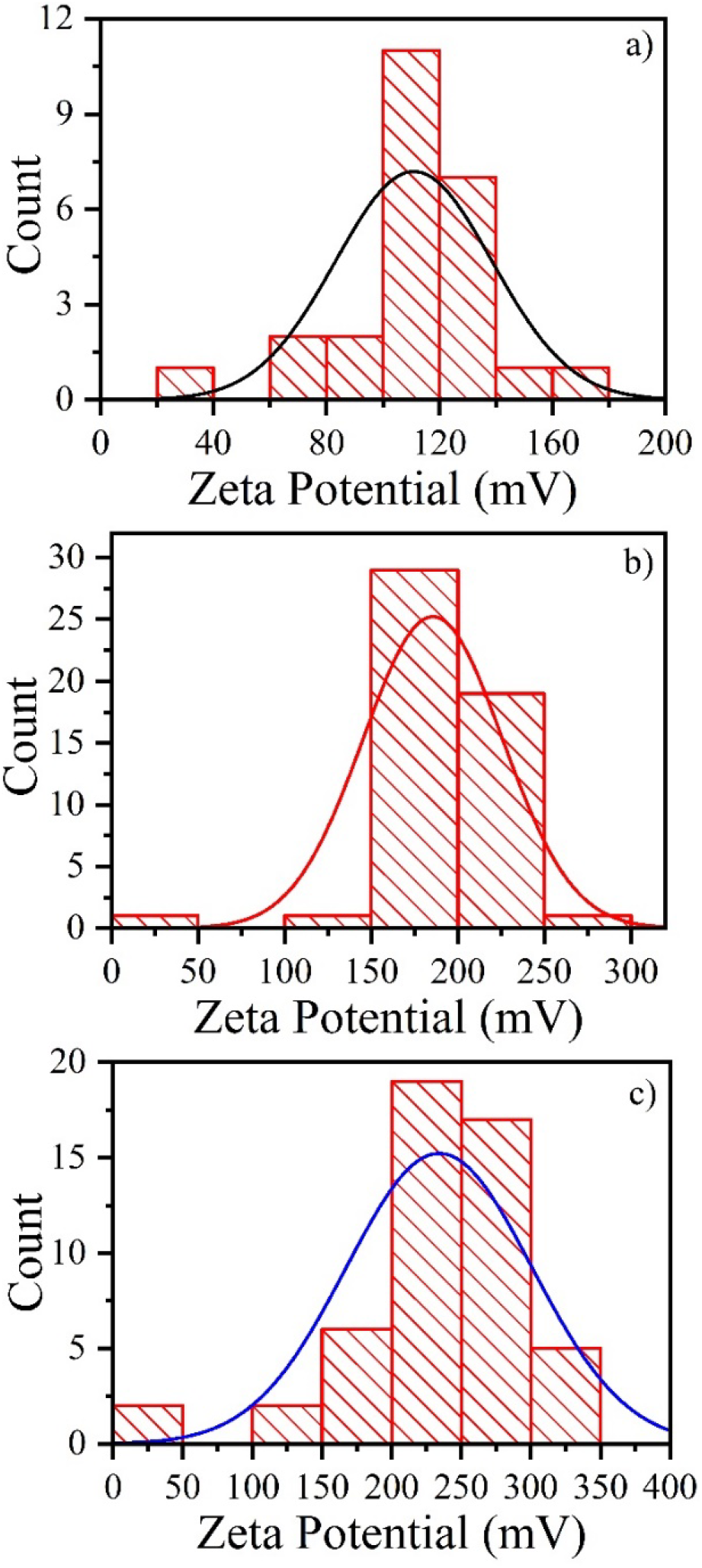
Zeta potential study of a) ZnO QDs, b) PVA-ZnO QDs, and c) PVP-ZnO QDs.

The selectivity study of ZnO QDs, PVA-ZnO QDs, and PVP-ZnO QDs towards ten antibiotics namely: Amoxicillin, Azaeothromycin, Ciprofloxacin, Floxacin, Gentamycin, Levofloxacin, Moxifloxacin, Norfloxacin, Tetracyclin, and Thiamphenocol was carried out. In the ZnO QDs, no specificity towards any antibiotic has been obtained. The peak wavelength in absorption study shows (fig 10(a)) no change with the interaction of antibiotics. There was a decrease in the fluorescence intensity of ZnO QDs when interacted with all the antibiotics. Therefore, no specificity has been achieved as shown in fig.10 (b).

**Fig. 10:**
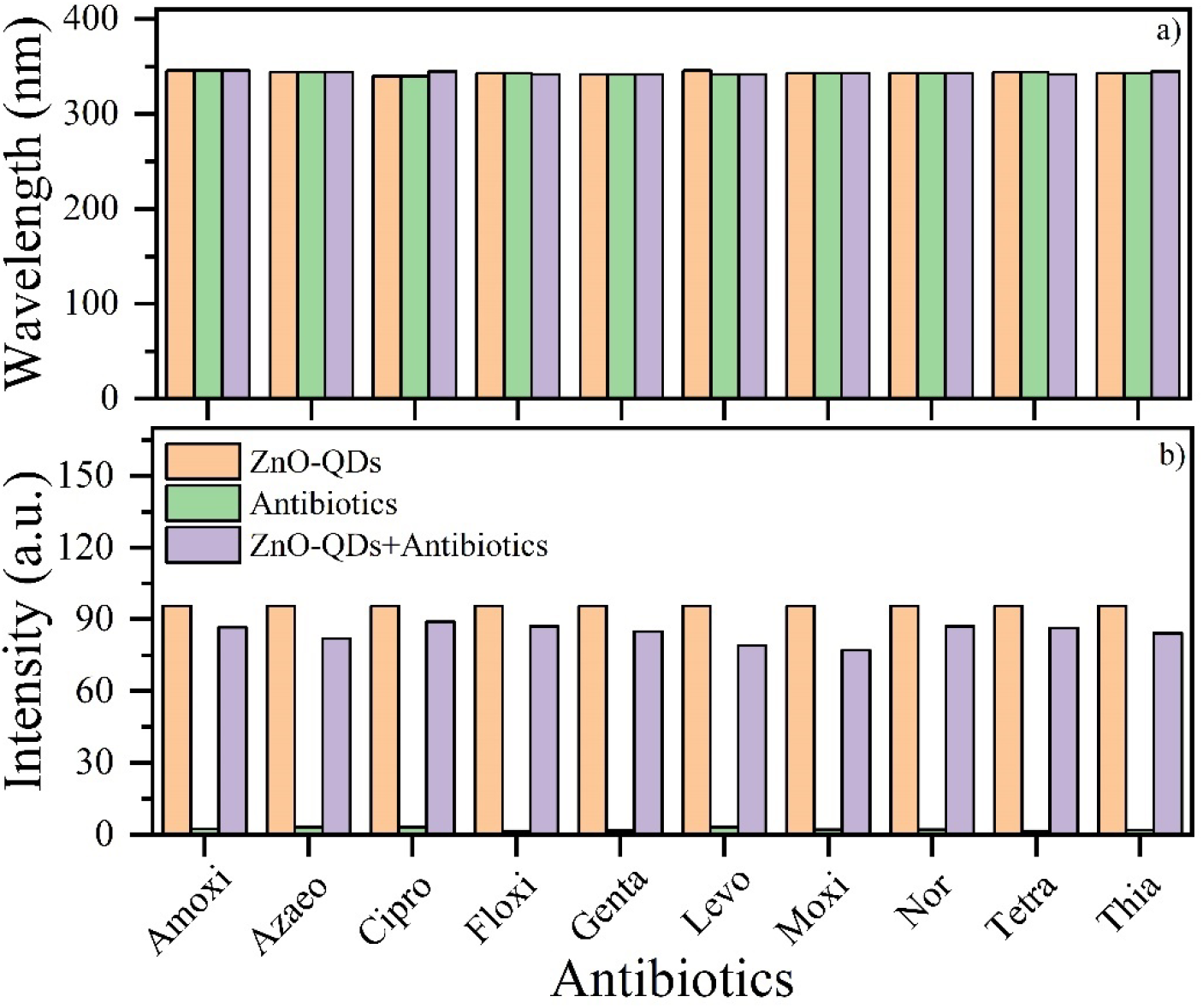
Selectivity study of ZnO QDs with various antibiotics a) peak wavelength change and b) fluorescence Intensity change

When the same study was performed with PVA-ZnO QDs, specificity towards ciprofloxacin has been obtained. The absorption peak wavelength study shows slight change in the wavelength with the interaction of ciprofloxacin as shown in fig 11(a). At the same time, the fluorescence intensity of both ZnO QDs and Cipro were decreased (fig 11(b)). And it was observed a mixed response of PVA-ZnO QDs when interacted with all the other antibiotics in the form of both the incarese and decraese in the intensity. No change in the wavelength of other antibiotics can be seen (fig 11(a)) which shows the specificity towards ciprolfoxacin for PVA-ZnO QDs.

**Fig. 11:**
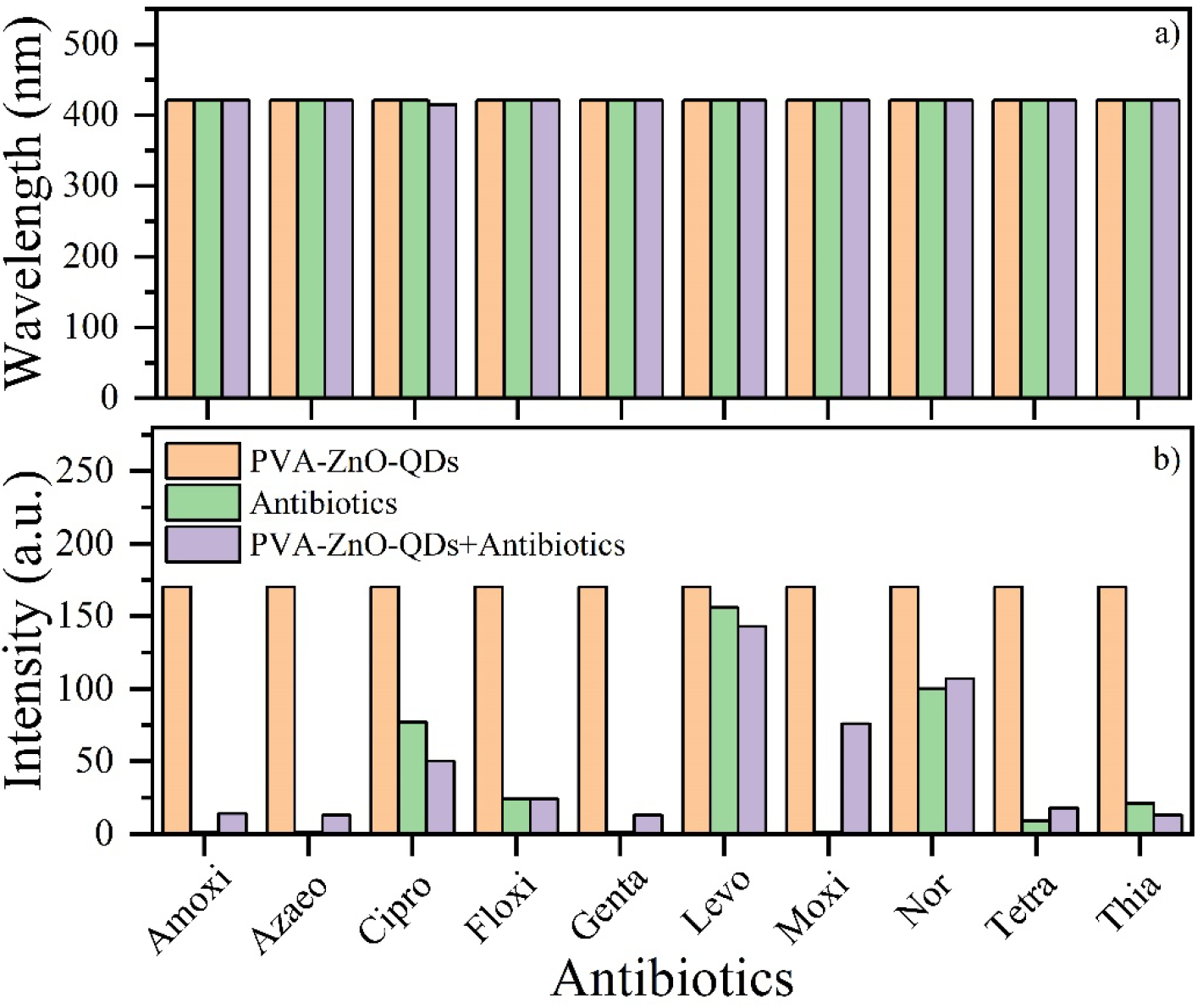
Selectivity study of PVA-ZnO QDs with various antibiotics a) peak wavelength change and b) fluorescence Intensity change

In the PVP-ZnO QDs, specificity towards moxifloxacin has been obtained. The absorption peak wavelength study shows a slight change in the peak wavelength when interacted with moxifloxacin as shown in figure 12(a). On the other hand, the fluorescence intensity of ZnO QDs was decreased after the interaction with Moxi. There has been mixed response of PVP-ZnO QDs when interacted with all the other antibiotics in the form of both the incarese and decraese in the intensity. No change in the wavelength of other antibiotics can be seen which shows the high specificity towards moxifloxacin for PVP-ZnO QDs as shown in fig. 12(b)).

**Fig. 12:**
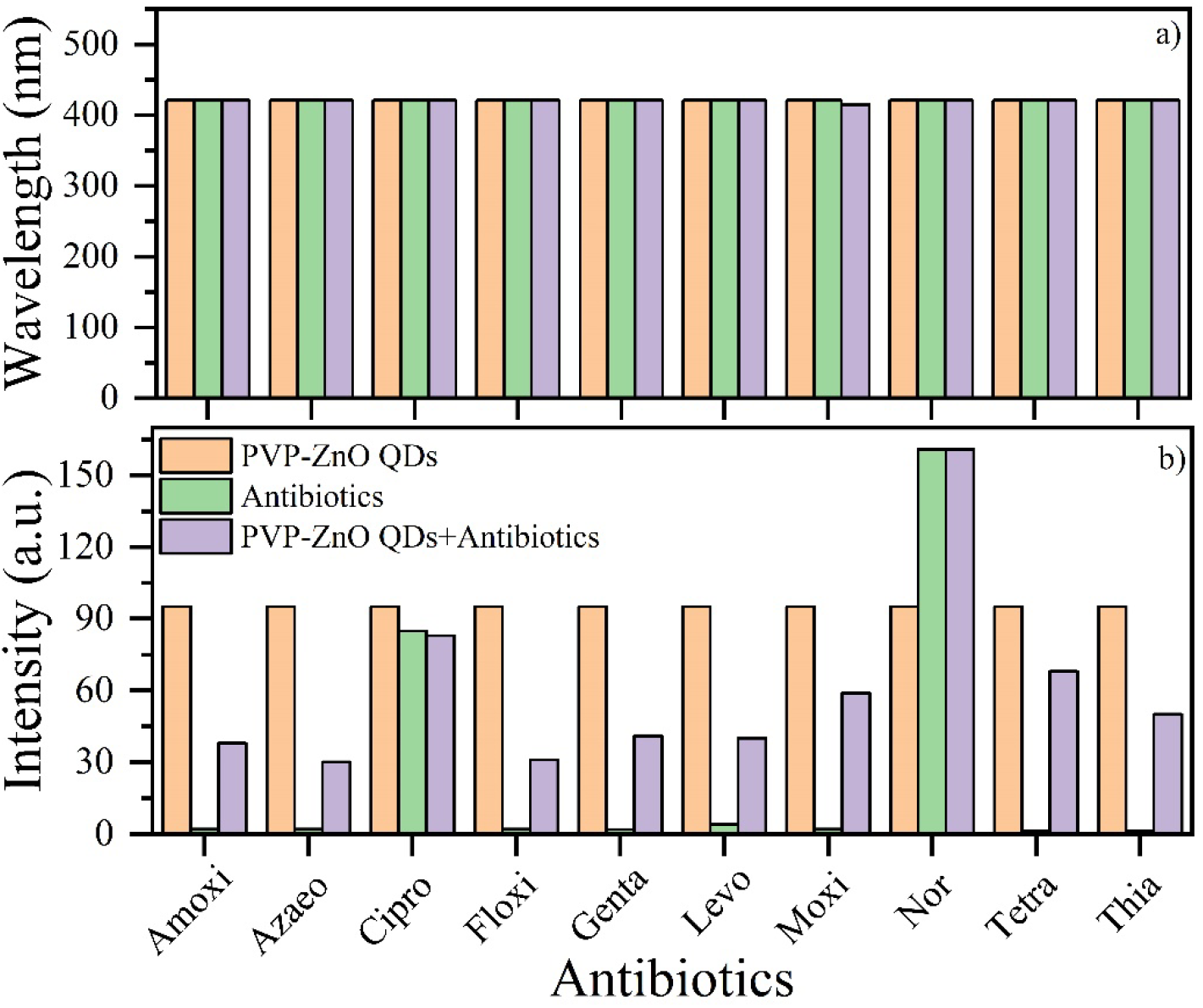
Selectivity study of PVP-ZnO QDs with various antibiotics a) peak wavelength change and b) fluorescence Intensity change

### 3.6 Ciprofloxacin sensing study against PVA-ZnO QDs

The sensing study of PVA-ZnO QDs towards ciprofloxacin, was performed by fluorescence study in the concentration range of 1 nM – 1 mM of ciprofloxacin as shown in fig 13. It is clear from the figure 13 that, complete quenching of the fluorescence of PVA-ZnO QDs and Cipro mixture obtained which shows the systematic interaction between the Cipro and PVA-ZnO QDs. The fluorescence peak intensity at each concentration was plotted against log of concentration of Cipro and a linear plot was obtained with two slopes as shown in inset of fig 13. The linearity ranges obtained were of 1 nM to 1 µM with R^2^=0.965 and 5 µM to 1 mM with R^2^=0.983.

**Fig.13:**
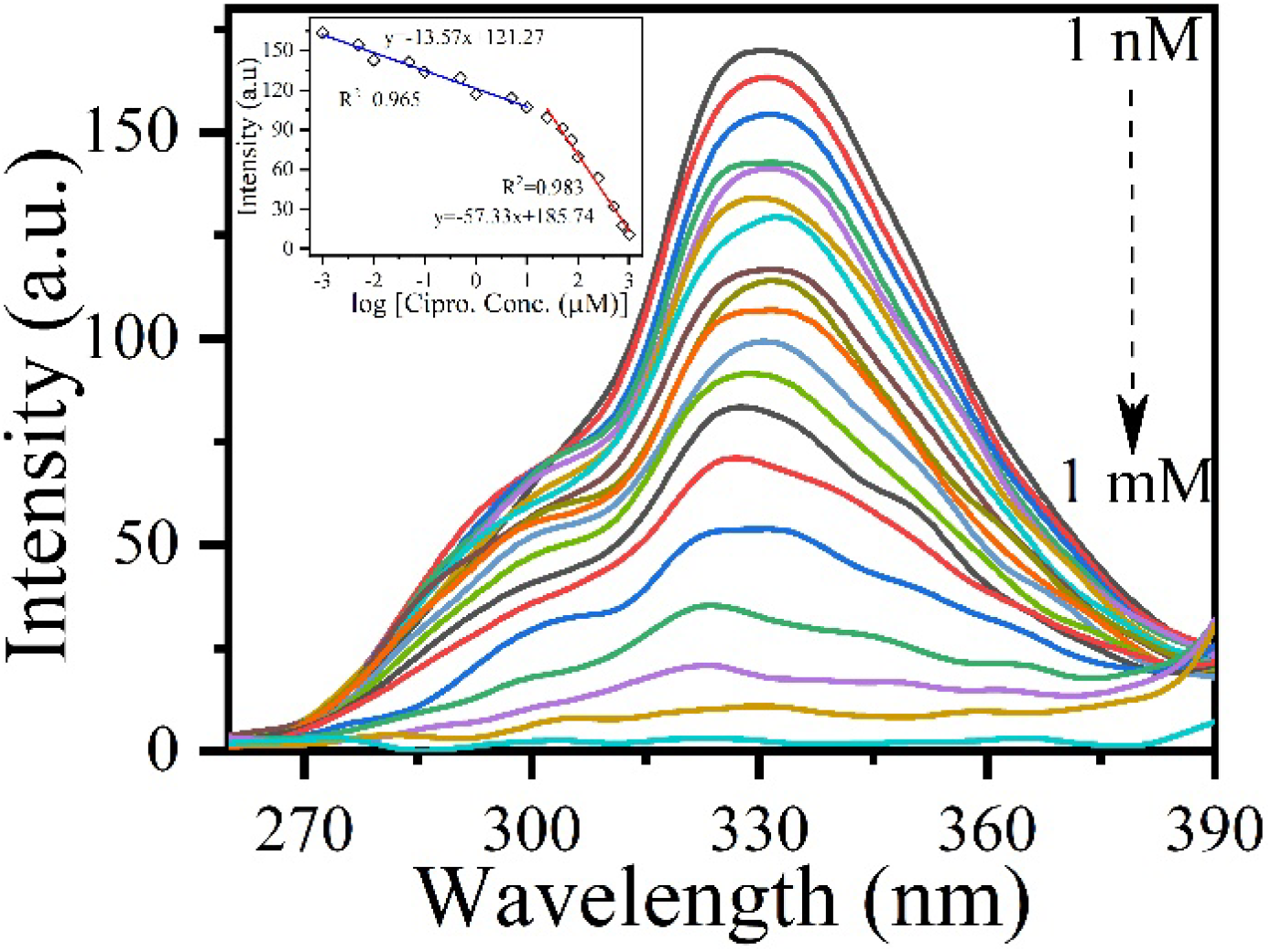
Sensing response study of PVA-ZnO QDs with ciprofloxacin

### 3.7. Ciprofloxacin spike study in tap water

Tap water was spiked with ciprofloxacin in the concentration range of 1 nM – 1 mM and perfomed the sensing with PVA-ZnO QDs using fluorescence spectra as shown in fig 14. It is clearly observed from fig 14 that, complete quenching of the fluorescence has been obtained as in the case of response study. The calibration curve was plotted in the spiked sample study also and shown in the inset of fig 14. Here also, two slopes obtained in the ranges of 1 nm to 1 µM with R^2^=0.98 and 5 µM to 1 mM with R^2^=0.99 as in the case of response study. These results indicate that the present system can be applicable for the sensing of Cipro in real samples also as contaminated water sources, hospital waste water, aquaculture sources etc.

**Fig.14:**
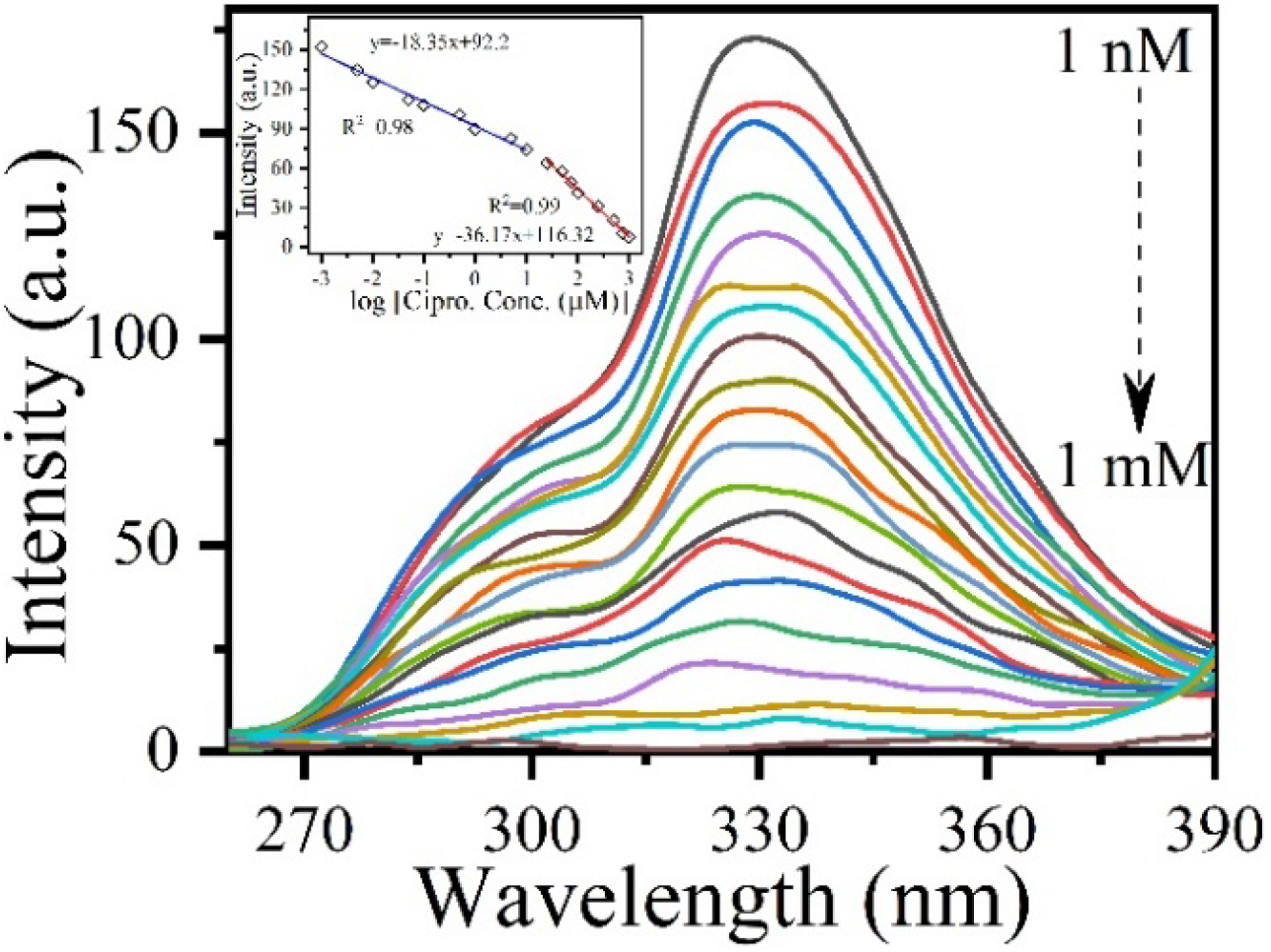
Spike study of PVA-ZnO QDs with ciprofloxacin antibiotic in the range of 1 nM to 1 mM in tap water.

### 3.8. Interference study-ciprofloxacin

Fig.15 shows the interference study performed on common interferents such as citric acid, ascorbic acid, cholesterol, aspartic acid, uric acid, folic acid, caseine, urea, glucose, sodium (Na^+^), potassium (K^2+^), calcium (Ca^2+^), copper (Cu^2+^), magnesium (Mg^2+^), manganese (Mn^2+^), and Zinc (Zn^2+^) that present in tap water and milk samples. Least change in the intensity can be seen for the Ciprofloxacin which confirms highly selective nature of the PVA-ZnO QDs towards ciprofloxacin.

**Fig.15:**
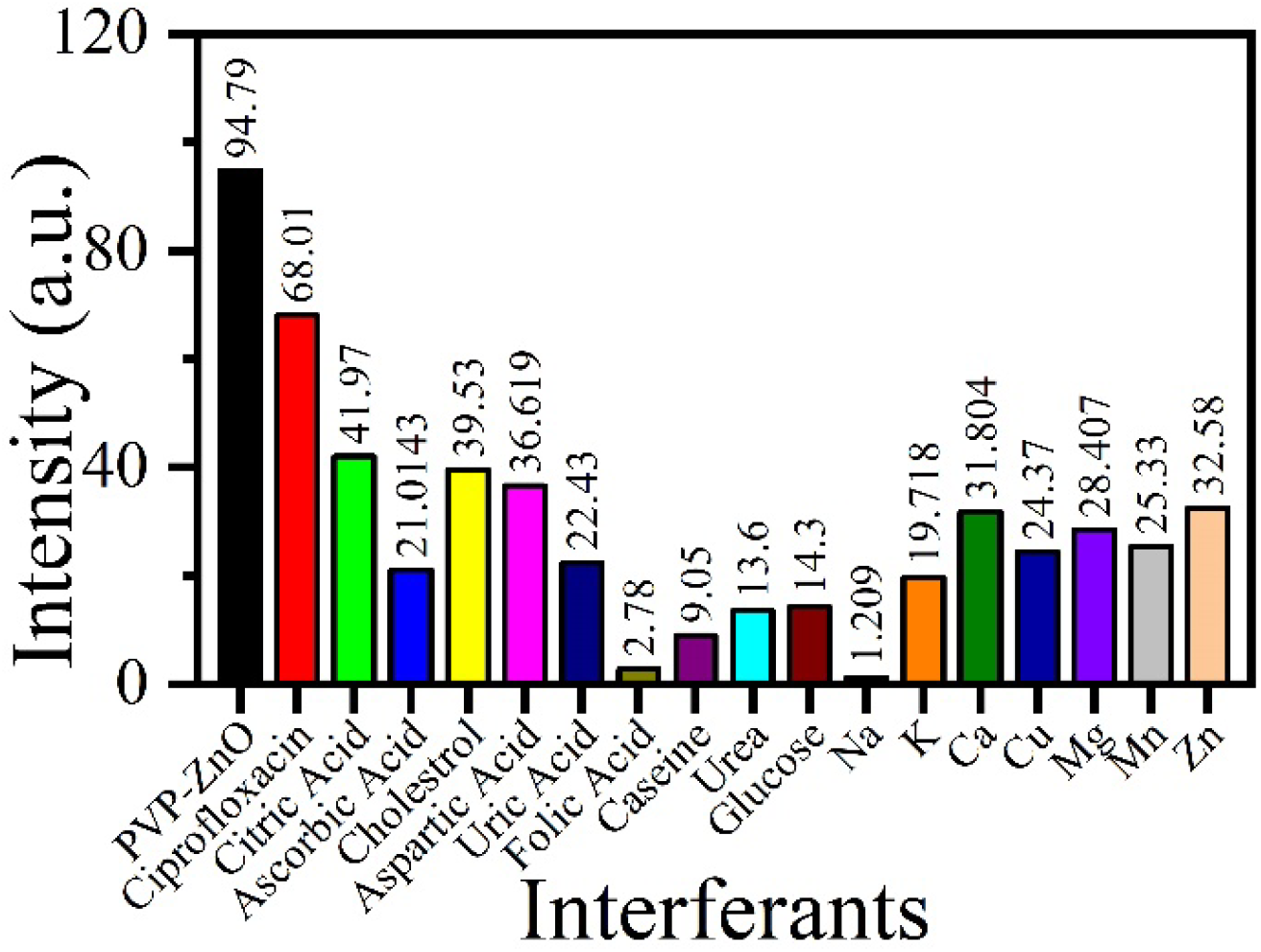
Interference study against multiple common interferants for ciprofloxacin interaction against PVA-ZnO QDs.

### 3.9. Moxifloxacin sensing study against PVP-ZnO QDs

The antibiotic sensing study was performed using fluorescence of PVP-ZnO QDs towards moxifloxacin, in the concentration range of 1 nM – 1 mM of Moxi as shown in fig 16. The fluorescence intensity was completely quenched at higher concentrations showing the interaction between the moxifloxacin and PVP-ZnO QDs. The calibration plot between the peak intensity and log of concentration, showed two slopes in the range of 1 nm to 1 µM with R^2^=0.956 and 5 µM to 1 mM with R^2^=0.99 as shown in inset of fig 16.

**Fig. 16:**
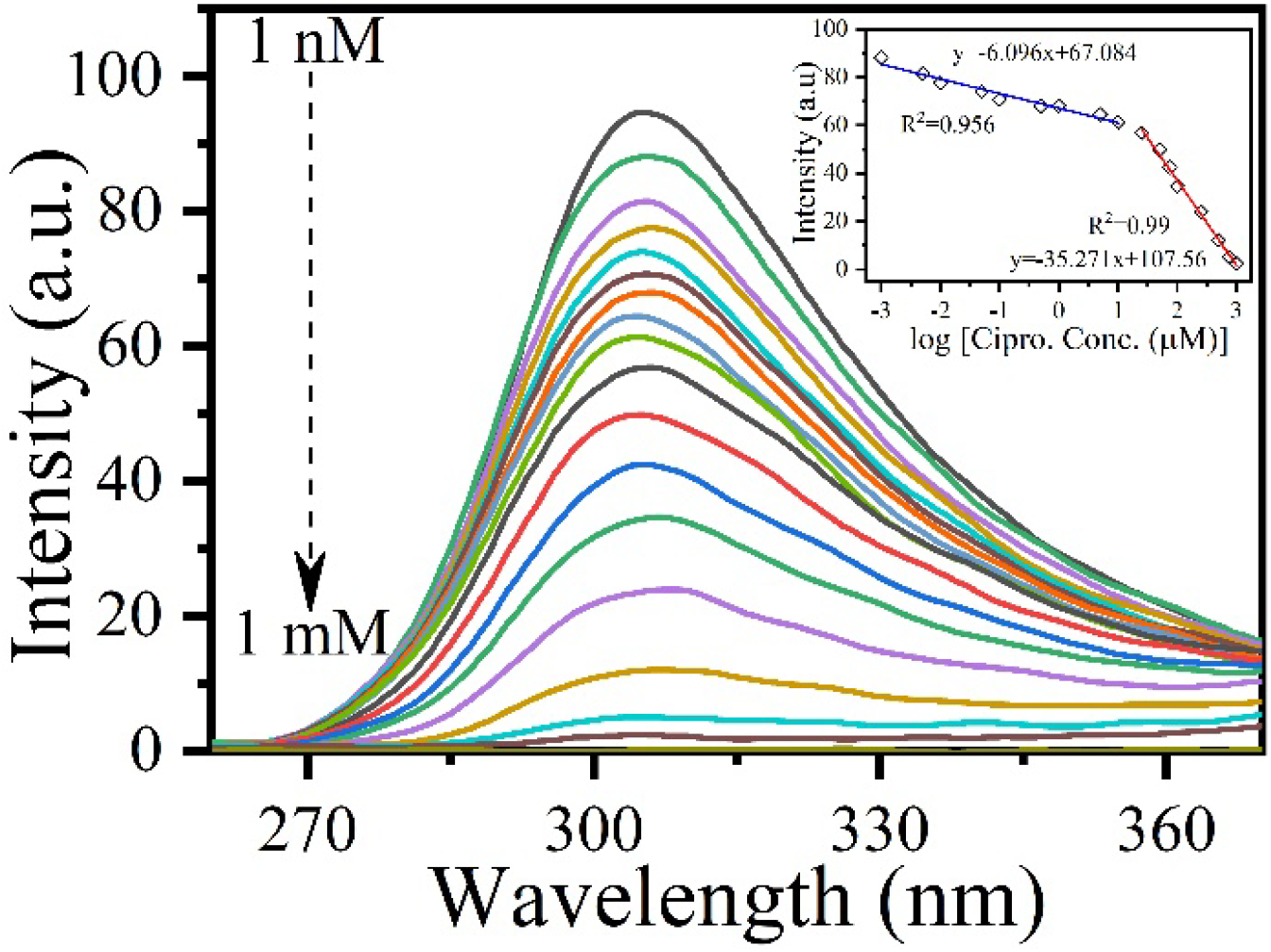
Response study of PVP-ZnO QDs with moxifloxacin

### 3.10 Moxifloxacin spike study in tap water

The spike study of PVP-ZnO QDs towards moxifloxacin, was also carried out in tap water using fluorescence spectra, by spiking the tap water with different concentrations of Moxi in the range of 1 nM – 1 mM and plotted in fig 17. It clearly shows the complete quenching of fluorescence as that of the response study showing the applicability of present system in real water samples. As shown in inset of fig 17, the calibration plot showed two slopes in the range of 1 nm to 1 µM with R^2^=0.956 and 5 µM to 1 mM with R^2^=0.98.

**Fig.17:**
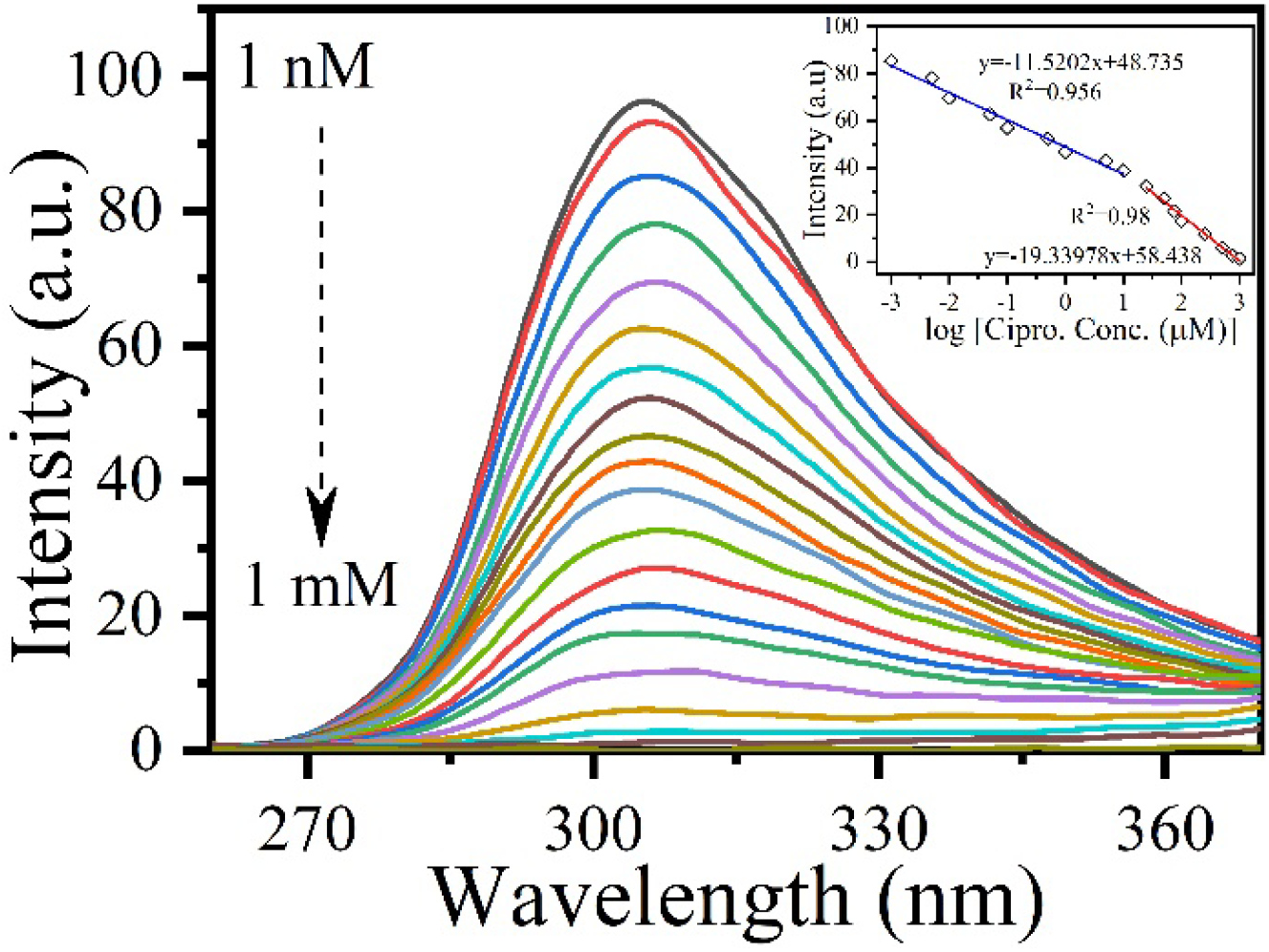
Spike study of PVP-ZnO QDs with moxifloxacin antibiotic in tap water

**Fig.17:**
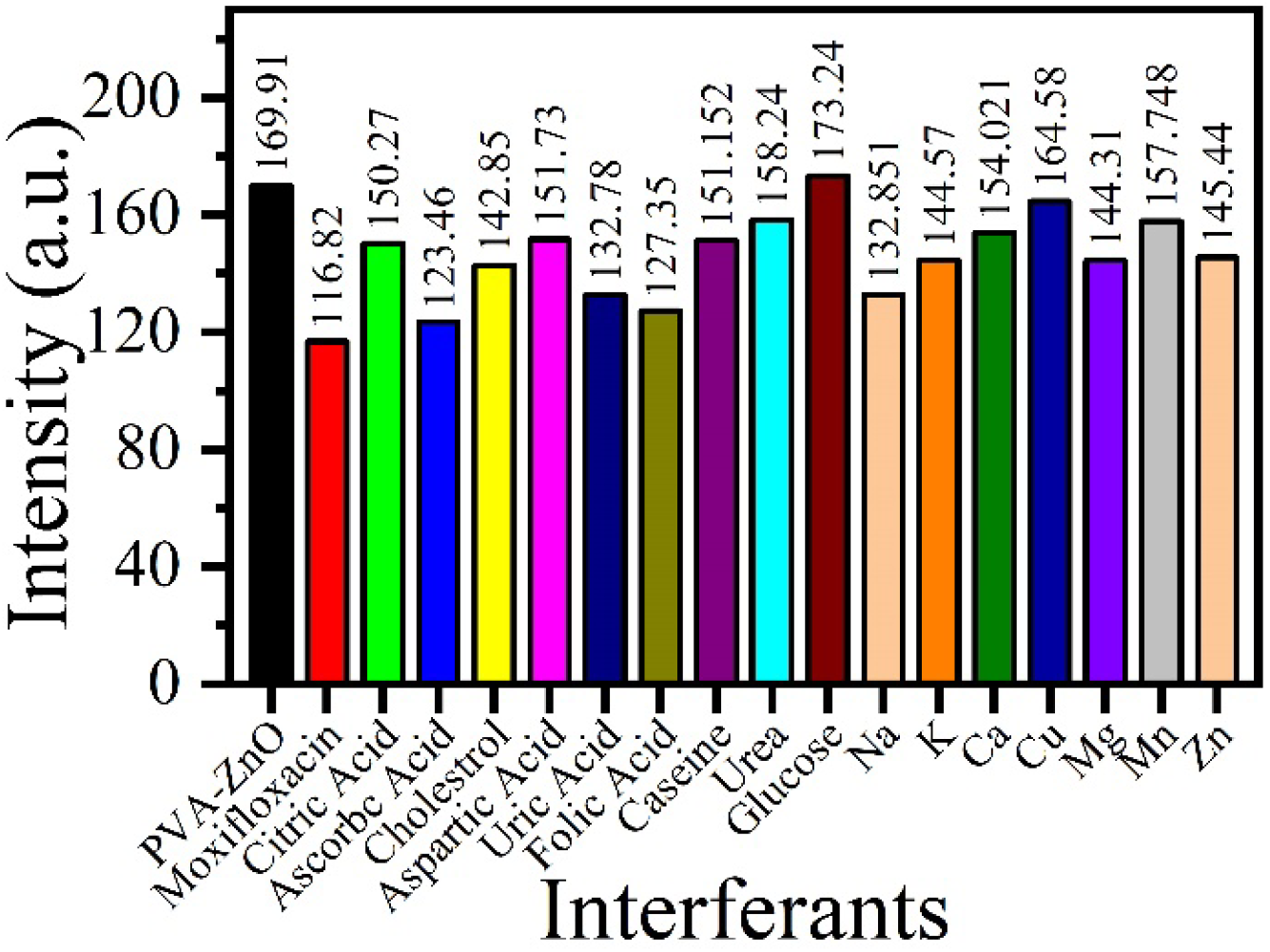
Interference study against common interferants for moxifloxacin

### 3.11 Interference study-moxifloxacin

Fig.18 shows the interference study performed on common interferents such as citric acid, ascorbic acid, cholesterol, aspartic acid, uric acid, folic acid, caseine, urea, glucose, sodium (Na^+^), potassium (K^2+^), calcium (Ca^2+^), copper (Cu^2+^), magnesium (Mg^2+^), manganese (Mn^2+^), and Zinc (Zn^2+^). Highest change in the intensity can be seen for the moxifloxacin which confirms highly selective nature of the PVP-ZnO QDs towards moxifloxacin.

**Fig.18:**
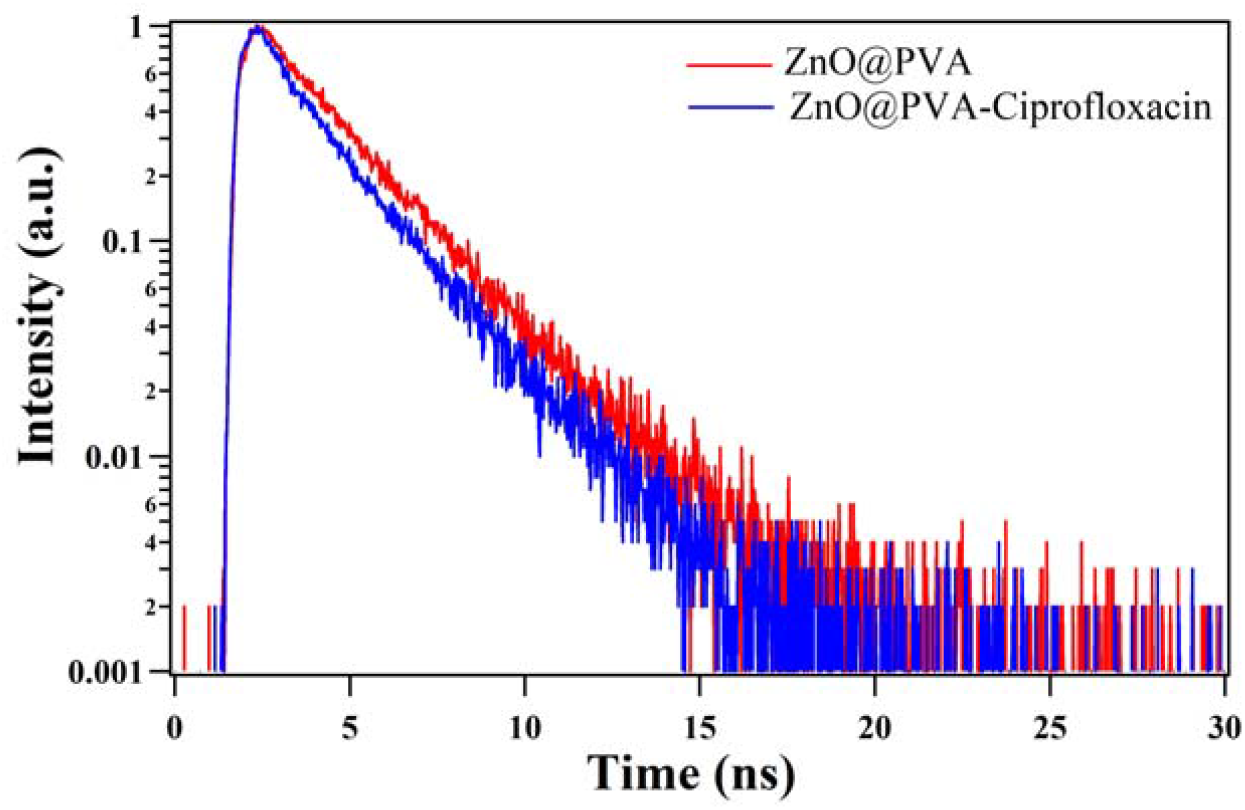
Fluorescence decay plot of PVA-ZnO QDs in presence and absence of Ciprofloxacin

### 3.12 Time-Resolved Fluorescence Spectroscopic analysis

The time-resolved fluorescence spectra of PVA-ZnO were recorded using time-correlated single-photon counting (TCSPC) instrument at 360 nm emission wavelength in the presence and absence of Ciprofloxacin. The measured data showed decrease in lifetime of ZnO-PVA in presence of Ciprofloxacin. These results indicated that Ciprofloxacin is quenching the ZnO-PVA QDs by dynamic quenching process. The decay plot of PVA-ZnO QDs in presence of Ciprofloxacin is plotted in the fig.18.

Whereas in case of PVP-ZnO QDs, the lifetime (from TCSPC study) did not show any change in the presence and absence of Moxifloxacin indicating the quenching is static in nature. The decay plot of PVP-ZnO QDs in presence of Moxifloxacin is plotted in the fig.19.

**Fig. 19:**
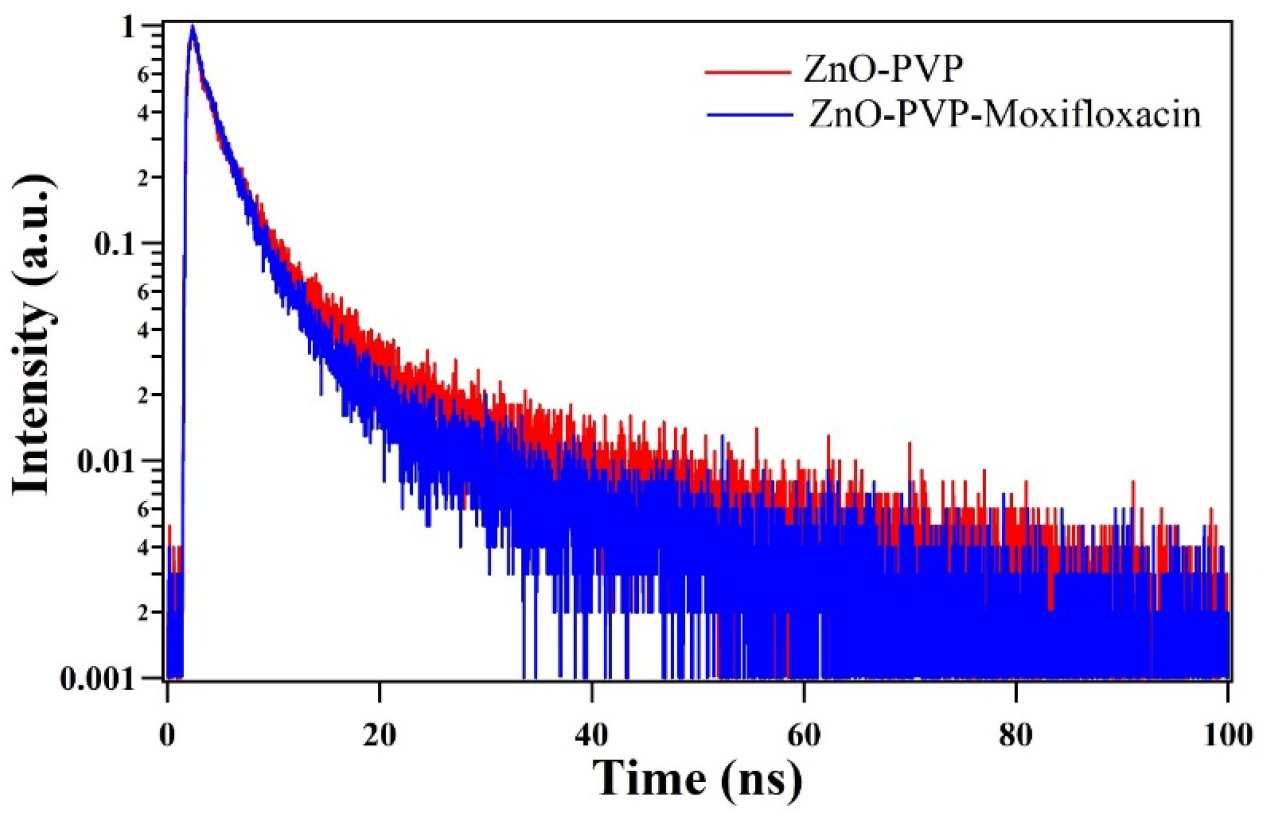
Fluorescence decay plot of PVP-ZnO QDs in presence and absence of Moxifloxacin

## 4. Conclusion

In the present study it is proved that, the polymer functionalization of ZnO QDs can tune the optical absorption and bandgap when functionalized with polymers like PVP and PVA. The Zeta potential studies reveal that the surface charge increased due to the polymer functionalization and it is observed that each of the sample carries positive charge. It is observed that the polymer functionalized ZnO QDs interacted specifically with different antibiotics such as PVA-ZnO QDs towards ciprofloxacin and PVP-ZnO QDs towards moxifloxacin. Whereas, the non-functionalised ZnO QDs did not show specifity. The fluoresecnce quenching of PVA-ZnO QDs by ciprofloxacin was dynamic in nature and PVP-ZnO QDs by moxifloxacin was in static nature. The spike sample studies in tap water showed promissing results towards applicability of the developed sensing systems in real samples. The specificity was proved by intereference studies.

## Author Contribution

AKV performed all the experimental, analyzed data and wrote the entire draft. TKD helped in experiment and data plotting. GBVS helped in analysis, editing and reviewed the draft. PS designed the research, provided the facility and edited the paper for final version.

---

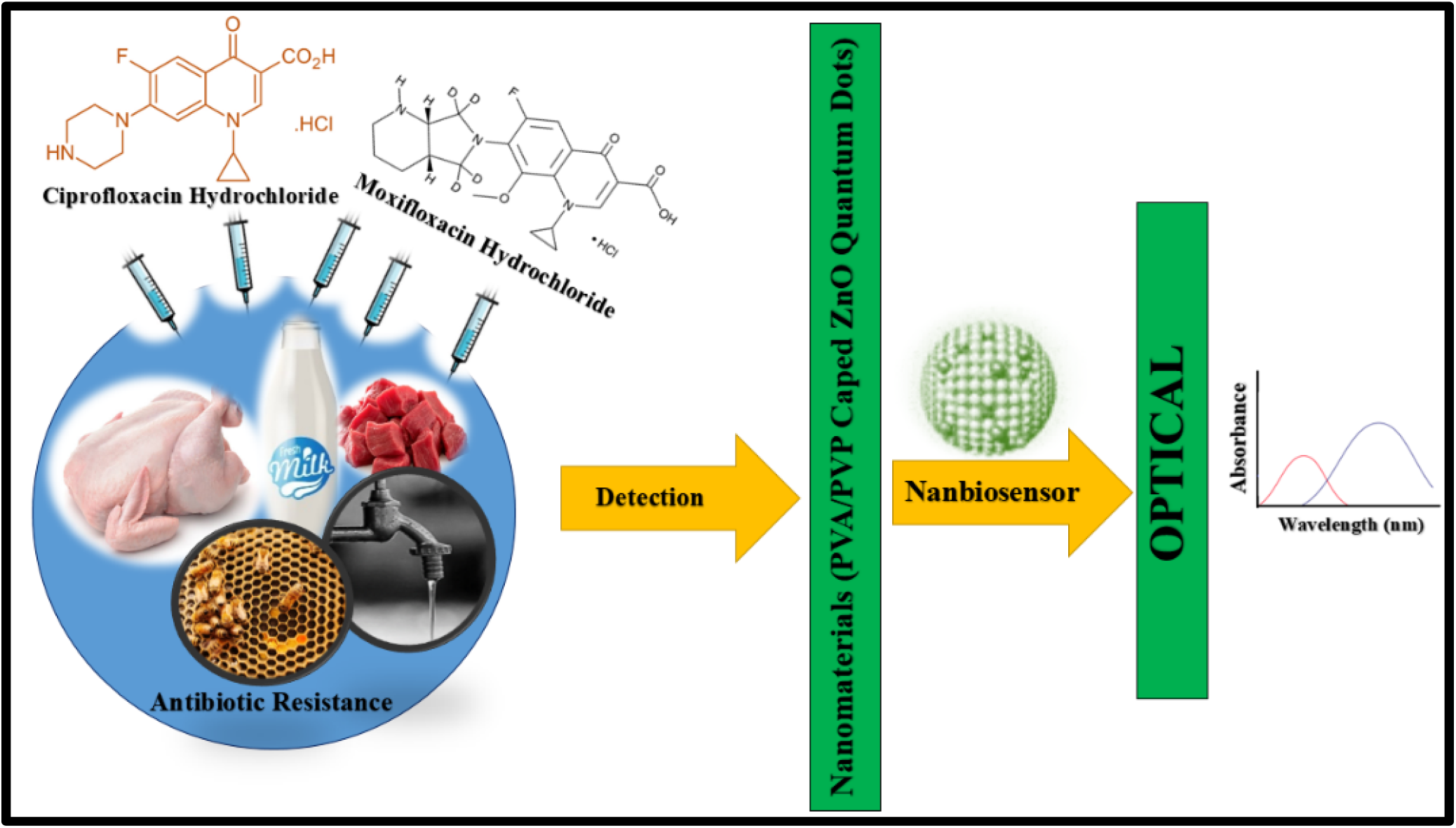

